# Detergent-Free Decellularization Preserves the Structural and Biological Integrity of Murine Tendon

**DOI:** 10.1101/2022.07.11.499615

**Authors:** Jason C. Marvin, Ai Mochida, Juan Paredes, Brenna Vaughn, Nelly Andarawis-Puri

## Abstract

Tissue decellularization has demonstrated widespread applications across numerous organ systems for tissue engineering and regenerative medicine applications. Decellularized tissues are expected to retain structural and/or compositional features of the natural extracellular matrix (ECM), enabling investigation of biochemical factors and cell-ECM interactions that drive tissue homeostasis, healing, and disease. However, the dense collagenous tendon matrix has limited the efficacy of traditional decellularization strategies without the aid of harsh chemical detergents and/or physical agitation that disrupt tissue integrity and denature proteins involved in regulating cell behavior. Here, we adapted and established the advantages of a detergent-free decellularization method that relies on Latrunculin B actin destabilization, alternating hypertonic-hypotonic salt and water incubations, nuclease-assisted elimination of cellular material, and protease inhibitor supplementation under aseptic conditions. Compared with previous tendon decellularization studies, our method minimized collagen denaturation while adequately removing cells and preserving bulk tissue alignment and mechanical properties. Furthermore, we demonstrated that decellularized tendon ECM-derived coatings isolated from different mouse strains, injury states (i.e., naïve and acutely injured/’provisional’), and anatomical sites harness distinct biochemical cues and robustly maintain tendon cell viability *in vitro*. Together, our work provides a simple and scalable decellularization method to facilitate mechanistic studies that will expand our fundamental understanding of tendon ECM and cell biology.

**Impact Statement:** In this study, we present a decellularization method for tendon that does not rely on any detergents or physical processing techniques. We assessed the impact of detergent-free decellularization using tissue, cellular, and molecular level analyses and validated the preservation of tendon structural organization, collagen molecular integrity, and ECM-associated biological cues that are essential for studying physiological cell-ECM interactions. Lastly, we demonstrated the success of this method on healthy and injured tendon environments, across mouse strains, and for different types of tendons, illustrating the utility of this approach for isolating the contributions of biochemical cues within unique tendon ECM microenvironments.

## Introduction

Tendinopathies, or tendon disorders, are debilitating conditions that account for over 30% of musculoskeletal consultations.^1^ The tendon extracellular matrix (ECM) is primarily composed of type I collagen fibers that are sparsely populated by resident tendon cells.^2^ Owing to their intrinsically limited healing capacity, ruptured tendons in adults typically heal by forming disorganized and mechanically impaired scar tissue (‘scar-mediated healing) that leads to long-term deficits in mobility, function, and quality of life.^3,4^ A gold standard for the treatment of tendinopathy is surgical reconstruction of the ruptured tendon ends using a donor graft (e.g., xenograft or patient-derived autografts and allografts). However, the success of these options has been restricted by challenges in the procurement of tendon grafts, post-operative retear rates as high as 94%^5,6^, and their limited capacity in attenuating further tissue degeneration.^2,4,7^ Despite the clinical prevalence of tendinopathies and our growing understanding of the hallmarks governing their pathogenesis, the biological mechanisms that ultimately drive effective repair and functional restoration of the tendon remain elusive.^8,9^ In particular, the role of local ECM microenvironmental cues in regulating tendon healing remains to be fully elucidated.

Tissue decellularization has emerged as a promising strategy to generate an acellular biologic scaffold to investigate the contributions of naturally-derived ECM constituents and topographical features in guiding cell behavior.^10,11^ Additionally, advances in methods for processing decellularized ECM (dECM) have given rise to several *in vitro* and *in vivo* applications, including 2D dECM-coated culture substrates^12,13^, injectable dECM-based biomaterials^14,15^, and 3D dECM-encapsulated hydrogels and electrospun scaffolds^12,15–19^ that retain tissue-specific and instructive cues that direct cell morphology, proliferation, and differentiation. However, the majority of tendon decellularization strategies employ harsh chemical detergents such as sodium dodecyl sulphate (SDS) and Triton X-100 and/or a combination of techniques that physically disrupt the ECM (e.g., repeated freeze-thaw cycles, ultrasonication, cryosectioning) that are known to negatively impact protein activity, tissue ultrastructural, and mechanical stability.^10,20–25^ Hwang *et* al. reported that even low concentrations of commonly used detergents can denature collagen molecules (i.e., irreversible unfolding of the collagen triple helix) in healthy porcine ligaments.^26^ Moreover, White *et al*. detected residual detergent fragments in decellularized tissues using time of flight secondary ion mass spectroscopy (ToF-SIMS) that were shown to disrupt the ECM surface ligand landscape and consequently led to adverse effects on reseeded cell behavior *in vitro*.^27^ Thus, there is mounting evidence in support of detergent-free decellularization approaches for tendon to more robustly recapitulate the native ECM biological environment.

In this study, we adapted and comprehensively characterized the effects of a detergent-free and non-proteolytic decellularization method that can be used to preserve the ECM integrity of mouse tendons across different genetic backgrounds, injury states, and anatomical sites. We hypothesized that tendons decellularized without detergents will be absent of any detectable molecular collagen denaturation or ECM destruction and therefore maintain their biological niche for cellular interrogation *in vitro*.

## Materials and Methods

### Animals and acute tendon injury model for tissue collection

All animal procedures were approved by the Cornell University Institutional Animal Care and Use Committee (IACUC). For primary cell isolations, male B6 mice at 16-to-18 weeks of age were bred in-house with the original dame and sire purchased from The Jackson Laboratory (#000664). For patellar tendon midsubstance punch surgeries, a subset of B6 mice were purchased directly from The Jackson Laboratory and acclimated for at least one week. To obtain a tendon biological environment distinct from that of B6 mice during injury, immunodeficient male NOD scid gamma (NSG) mice at 16-to-18 weeks of age were obtained from the Cornell University Progressive Assessment of Therapeutics (PATh) PDX Facility. Mice were housed with up to five animals per cage in an alternating 12-hour light/dark cycles with ab libitum access to chow and water.

To generate the injured/‘provisional’ tendon ECM, a 1-mm diameter, full-thickness biopsy punch was introduced into the left patellar tendon midsubstance of B6 and NSG mice as described previously.^28,29^ Briefly, mice were anesthetized with isoflurane (2% volume, 0.3 L/min) and received pre-operative buprenorphine (1 mg/kg body weight) via intraperitoneal administration. The fur at the surgical site was removed before making an incision directly above the knee cavity to expose the patellar tendon. A polyurethane-coated stainless-steel backing was inserted underneath the tendon before using a biopsy punch to create the injury defect. The backing was removed before closing the skin incision using 6-0 prolene sutures (Ethicon Inc., #8695G). Mice resumed cage activity. At 1-week post-injury, mice were euthanized via carbon dioxide (CO_2_) inhalation followed by cervical dislocation. Hole-punched (provisional ECM) and uninjured contralateral tendons (naïve ECM) were immediately harvested, flash-frozen in liquid nitrogen (LN_2_), and stored at −80°C until decellularization. To demonstrate the utility of this decellularization method for other tendon tissues, uninjured B6 Achilles tendons were also collected.

For evaluation of bulk tensile mechanical properties, the patella-tendon-tibia complex was harvested from both limbs of uninjured B6 mice, carefully cleaned of the surrounding musculature, flash-frozen in LN_2_, and stored at −80°C until decellularization.

### Primary mouse tendon cell isolation and culture

Cells were routinely cultured in basal media consisting of low-glucose Dulbecco’s modified Eagle’s medium (DMEM; Corning, #10-014) supplemented with 10% (v/v) lot-selected fetal bovine serum (FBS; Corning, #35-015-CV) and 1% (v/v) penicillin-streptomycin-amphotericin B (PSA; Thermo Fisher Scientific, #15240062). All experiments were conducted at passage 3 with media replenished every 2 days. All cells lines used in this study were negative for mycoplasma contamination (Lonza, #LT07-703).

B6 patellar tendon cells were isolated from uninjured animals as described previously.^29^ After euthanizing animals, the patellar tendon was harvested from both limbs from a total of five mice, stripped of the sheath and fat pad, and digested with a mixture of collagenase type I (2 mg/mL of digest solution) and collagenase type IV (1 mg/mL of digest solution) in serum-free DMEM for up to 2 hours at 37°C on a rocking shaker. Single-cell suspensions were obtained by passing tissue digests through a 70-μm strainer before first seeding cells onto a tissue culture-treated flask (passage 0).

### Decellularization of tendon tissue and preparation of ECM-coated substrates

Naïve and provisional mouse tendons were decellularized using an adapted detergent-free decellularization method^29,30^ for skeletal muscle^31^ based on Latrunculin A (Biovision, #2182-1) and alternating hypertonic-hypotonic solutions. Incubations steps were performed with 1 mL of solution (2 mL for mechanical testing samples to ensure that the tissue was fully submerged) in 2.0-mL microcentrifuge tubes (Corning, #MCT-150-C-S) with agitation at 450 rotations per minute (RPM) using an orbital shaker. All solutions with the exception of Latrunculin B were utilized at room temperature and supplemented with a 1X protease inhibitor cocktail (Thermo Fisher Scientific, #78425) to minimize proteolysis. To limit the risk of contamination, tissue handling and solution changes were done in a biosafety cabinet using aseptic technique. Lastly, salt solutions were prepared fresh in sterile DI water for each decellularization batch.

First, frozen tendons were thawed and individually incubated with 50 nM Latrunculin B (Biovision, #2182-1) for 2 hours at 37°C to disrupt actin polymerization. Second, tissue samples were then washed once with deionized (DI) water for 30 minutes followed by 0.6 M potassium chloride (KCl) for 2 hours, another DI water wash step for 30 minutes, and finally 1.0 M potassium iodide (KI) for 2 hours to induce osmolysis. Third, samples were then washed in DI water for 12 hours, subjected to the same alternating hypertonic-hypotonic solutions, incubated with 1X phosphate-buffered saline (PBS) supplemented with 1 kU/mL of Pierce Universal Nuclease (Thermo Fisher Scientific, #88702) for 2 hours to remove cellular debris, and then finally washed with 1X PBS for 48 hours with the solution changed each day. All samples were stored at 20°C until use.

For biochemical analyses, decellularized tendons were lyophilized for 72 hours, individually weighed to obtain the dry mass, and then incubated with 200 µL of papain digest solution (0.2 mg/mL; Sigma-Aldrich, #P4762) in 0.5-mL microcentrifuge tubes (Thermo Fisher Scientific, #AM12300). Next, samples were vortexed to sediment the tissue to the bottom of the tube before digesting for 16-to-18 hours at 65°C inside a tube rack on a rocking shaker until completely digested. Papain digests were stored at −20°C until analysis.

To make decellularized ECM-coated substrates, lyophilized tendons were first pooled up to 10 mg (dry weight) inside 2.0-mL low-binding tubes with screw caps and O-rings (Omni, Inc., #19-660-1000). Next, samples were mechanically homogenized (Omni, Inc., Bead Ruptor 24) using five 2.8-mm ceramic beads (Omni, Inc., #19-646) per tube for five pulverization cycles (6.0 meters/second velocity for 15 seconds per cycle) with flash-freezing in LN_2_ between each cycle. Samples were then solubilized (10 mg of dry tissue/mL of solution) with pepsin (1 mg/mL; Sigma-Aldrich, #P7012) in 0.1 M hydrochloric acid (HCl) for 72 hours at 4°C at 150 RPM, diluted to a working concentration of 1 mg of dry tissue/mL solution in 0.1 M acetic acid (AcOH), and stored at −80°C until use. Coatings were prepared by adding 50 µL of solubilized ECM to flat-bottom 96-well plates overnight at 4°C which were subsequently washed twice with sterile 1X PBS immediately prior to seeding cells.

### Biochemical analysis of dsDNA and sulfated glycosaminoglycan (sGAG) content

Biochemical analyses were performed on non-decellularized and decellularized samples using the papain digests. To determine the efficacy of the decellularization method in removing cellular materials, total double-stranded DNA (dsDNA) content was measured using a commercial Quant-iT PicoGreen kit (Thermo Fisher Scientific, P11496) as per the manufacturer’s instructions. Fluorescence was measured at excitation/emission wavelengths of 480/520 on a SpectraMax i3X Multi-Mode Microplate Reader (Molecular Devices). To determine the impact of detergent-free decellularization on the preservation of non-collagenous ECM constituents, sGAG content was measured using the dimethylmethylene blue (DMMB) assay^32^. After the addition of the DMMB solution to the samples, absorbances were immediately measured at 540 and 595 nm to calculate the sample concentration by subtracting the absorbance at 595 nm from the absorbance at 540 nm following a blank offset of DI water only.

### Histology

Non-decellularized and decellularized B6 patellar tendons were embedded in optimal cutting temperature (OCT) compound and cryosectioned sagittally at a thickness of 6 µm. To determine the effect of decellularization on tissue structure and removal of cellular materials, cryosections were stained with hematoxylin and eosin (H&E) and 4′,6-diamidino-2-phenylindole (DAPI), respectively. Images were acquired at 40X magnification on an inverted brightfield microscope (H&E) or Axio Observer Z1 epifluorescence microscope (DAPI).

### Fluorescent assessment of collagen denaturation with CF-CHP

To determine the extent of collagen denaturation after detergent-free decellularization, non-decellularized and decellularized cryosections were stained with 20 µM of carboxyfluorescein-labeled collagen hybridizing peptide^33–35^ (CF-CHP; Echelon Biosciences, #FLU300) in 1X PBS as per the manufacturer’s instructions. Positive controls for CF-CHP staining were prepared by heating uninjured B6 patellar tendons in 1X PBS at 65°C for 1 minute and cryosectioning as described above. Trimeric CF-CHP solution was heated at 80°C for 5 minutes to obtain thermally-dissociated monomeric strands. Monomeric CF-CHP solution was incubated at −80 °C for 15 seconds and then quenched to room temperature. Cryosections were stained with 25 µL of CF-CHP, incubated at 4°C for 12 hours in a humidified chamber, and carefully washed thrice in 1X PBS for 5 minutes per wash to remove unbound CF-CHP. Lastly, stained cryosections were dehydrated using a graded alcohol series and mounted in EUKITT® medium (Electron Microscopy Sciences). Images were acquired on a Zeiss LSM 710 confocal microscope (Zeiss) at 10X magnification and 488 nm laser excitation. Identical imaging parameters were used for all samples. Integrated density measurements of CF-CHP staining indicative of denatured collagen were quantified in ImageJ by a blinded user using a custom MATLAB script to exclude sectioning artifacts from analysis. A total of 2-4 stained sections (spaced at least 100-150 μm apart from each other) taken throughout the full thickness of the tendon were averaged for each sample to account for any variation in the penetration of decellularization reagents.

### Bulk tensile testing

Non-decellularized and decellularized B6 patella-tendon-tibia complexes were secured using custom grips in a 1X PBS bath at room temperature. A nominal 0.15 N pre-load was applied and then followed by preconditioning for 15 cycles at 1% strain and a frequency of 1 Hz. Stress relaxation was assessed at 5% strain (5% strain/second) with a recovery time of 300 seconds. After stress relaxation testing, tendons were held for an additional 300 seconds under no load. Pre-conditioning was then re-applied as described above before pulling samples uniaxially to failure at a rate of 0.1% strain/second to record the ultimate load and stiffness. Tendon cross-sectional area was calculated from digital images taken using a monochrome industrial camera (DMK 33UX250) during the 0.15 N pre-load prior to mechanical testing.

### Assessment of Cytocompatibility using live-dead cell staining

To assess the cytocompatibility of biochemical cues derived from the decellularized ECM, B6 patellar tendon cells were seeded (7,500 cells/well) in basal media onto B6 naïve or provisional ECM-coated 96-well glass-bottom plates (Greiner Bio-One, #655892). Non-coated wells served as controls. After adhering for 4 hours, cells were rinsed once with plain DMEM and then cultured in FluoroBrite DMEM (Thermo Fisher Scientific, #A1896701) supplemented with 1% lot-selected FBS and 1% PSA for 3 days. Live/dead cell staining was performed by incubating cells with 5 µM calcein AM and 3 µM propidium iodide (PI) for 30 minutes. Images were then immediately acquired on a Zeiss Axio Observer Z1 epifluorescence microscope at 2.5X magnification. Identical imaging parameters were used for all samples. Cell viability was calculated as the percentage of calcein-positive cells divided by the total number of cells. Three technical replicates were averaged for each biological replicate.

### Analysis of single-cell morphology in vitro

To functionally evaluate if the altered provisional ECM composition of NSG tendons due to their suppressed immune response is preserved with decellularization, B6 patellar tendon cells were seeded (2,000 cells/well) in basal media onto NSG naïve or provisional ECM-coated 96-well glass-bottom plates for 18 hours. Cells were then fixed with 4% paraformaldehyde for 30 minutes, permeabilized with 0.1% Triton X-100 for 15 minutes, blocked with 2.5% normal horse serum (Vector Laboratories, #S-2012) for 30 minutes, and concurrently stained with Alexa Fluor 488 Phalloidin (1:400 dilution; Thermo Fisher Scientific, #A12379) and DAPI (1:1000 dilution) in 2.5% normal horse serum for 1 hour while covered from light. All immunostaining steps were performed at room temperature. Cells were then rinsed twice and mounted with 1X PBS before images were acquired on a Zeiss Axio Observer Z1 epifluorescence microscope at 20X magnification. Single-cell morphological parameters, including cell spreading area, cell perimeter, and circularity (calculated as 4*π*cell spreading area/perimeter^2^) were quantified in ImageJ by a blinded user.

### Statistical Analysis

Biochemical and mechanical analyses for non-decellularized and decellularized samples were compared using an unpaired Student’s t-test. CF-CHP integrated density and cell viability between uncoated control, naïve, and provisional ECM-coated substrates were compared using a one-way analysis of variance (ANOVA) with post-hoc Tukey. Single-cell morphology between naïve and provisional ECM-coated substrates was compared using a one-way Welch ANOVA with post-hoc Games-Howell. All experiments were conducted with a minimum of 3 biological replicates with each biological replicate representing a unique tendon sample or cell line.

## Results

### Detergent-free decellularization preserved tissue structure and removed cells

Supporting our hypothesis, histological examination of H&E-stained naïve B6 patellar tendons revealed the preservation of highly aligned collagen fibers characteristic of native tendon structure following decellularization (**Fig. 1A**; top panel). Both H&E and DAPI staining (**Fig. 1A**) illustrated the removal of resident cells from naïve tendons after decellularization. Corroborating these visual observations, dsDNA content in B6 naïve and provisional tendons was significantly reduced from 1039.14 ± 184.1 ng/mg and 2957.37 ± 298.1 ng/mg, respectively, to 47.64 ± 5.36 (*P* < 0.0001) ng/mg and 184.6 ± 24.32 ng/mg (*P* < 0.0001) after decellularization (**Fig. 1B**). Similarly, dsDNA content in B6 naïve Achilles tendons was significantly reduced from 425.50 ± 5.17 ng/mg to 26.61 ± 0.55 ng/mg after decellularization (data not shown), establishing the efficacy of this decellularization method for different tendon tissues. As expected, the effect of decellularization on reducing overall dsDNA content was significantly greater (*P* = 0.0041) in naïve (95.42% ± 0.52% reduction) compared to provisional tendons (93.76% ± 0.82% reduction), which is presumably attributed to the nearly three-fold increase in cellularity and formation of dense granular tissue (i.e., lower penetration of decellularization reagents) during acute tendon injury.

**Figure 1.**
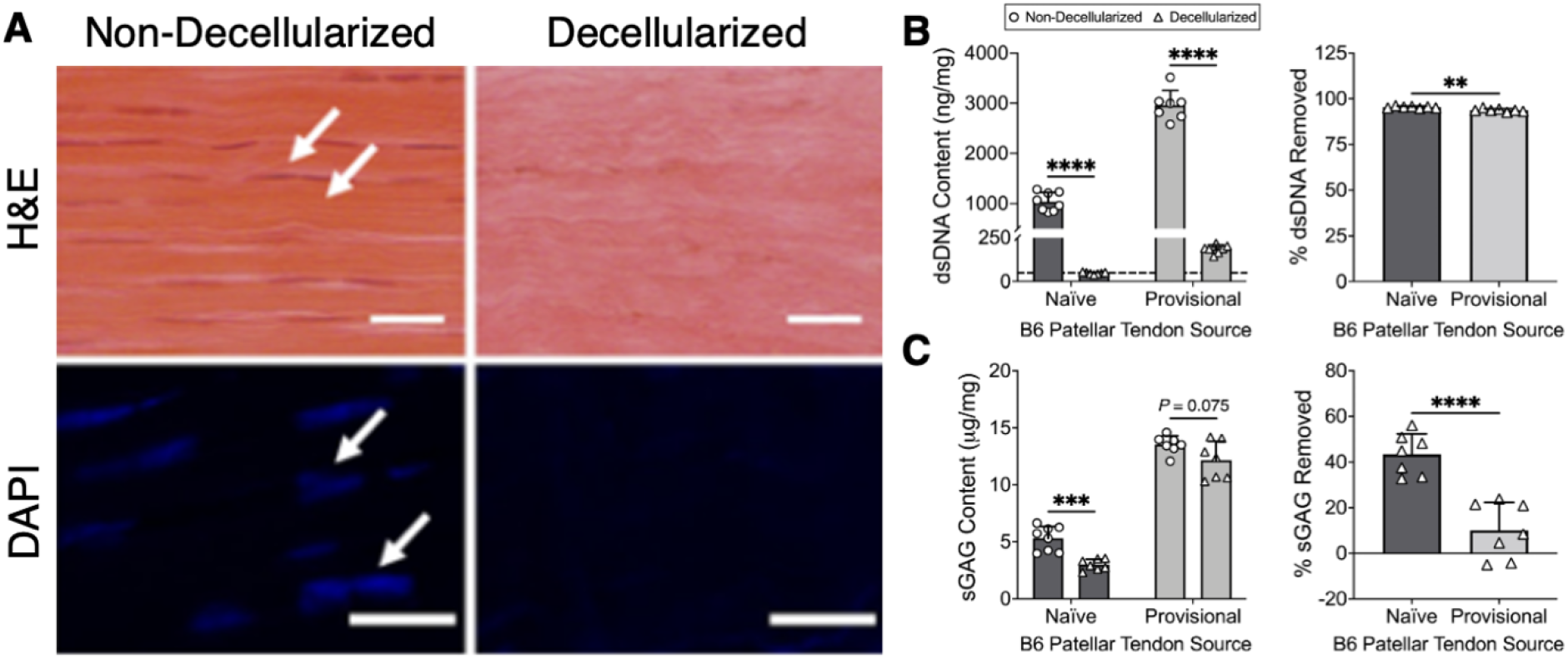
Histological and biochemical evaluation of non-decellularized and decellularized B6 patellar tendons. (**A**) Representative H&E (top panel; scale bar, 100 μm) and DAPI staining (bottom panel; scale bar, 25 μm) demonstrated preservation of collagen fiber alignment while removing cellular material. White arrows indicate cells. *N* = 6 samples per group. (**B**) Total dsDNA and (**C**) sGAG content in B6 naïve and provisional tendons were reduced before and after decellularization as measured by the Quant-iT PicoGreen and DMMB assays, respectively. *N* = 7-8 samples per group. Dashed black line indicates 50 ng/mg threshold for dsDNA content. Data is presented as mean ± standard deviation. \**\*P* < 0.01, \*\*\**P* < 0.001, \*\*\*\**P* < 0.0001.

The sGAG content in B6 naïve and provisional tendons was reduced from 5.29 ± 1.08 µg/mg and 13.53 ± 0.79 µg/mg, respectively, to 3.00 ± 0.48 µg/mg (*P* = 0.0002) and 12.17 ± 1.66 µg/mg (*P* = 0.075) after decellularization (**Fig. 1C**). Interestingly, the overall reduction in sGAG content was significantly lower (*P* < 0.0001) in provisional (10.01% ± 12.25%) compared to naïve (43.38% ± 9.03%). These results indicate that the sGAG-rich provisional ECM composition produced during the early proliferative phase of wound healing^36,37^ is largely maintained following detergent-free decellularization.

### Detergent-free decellularized tendons showed no signs of collagen denaturation

Collagen denaturation was not visually detected in non-decellularized and decellularized B6 naïve patellar tendons as evidenced by a lack of CF-CHP staining compared to the heat denatured control group (**Fig. 2A**). Furthermore, there was no difference in CF-CHP staining intensity between superficial and deeper cryosections of the tendon. Supporting our hypothesis, CF-CHP integrated density of non-decellularized and decellularized samples were comparable and both significantly lower (*P* < 0.0001 for both) than that of the heat denatured control group (**Fig. 2 B**), confirming that detergent-free decellularization combined with protease inhibitors limited proteolytic degradation of the tendon ECM environment.

**Figure 2.**
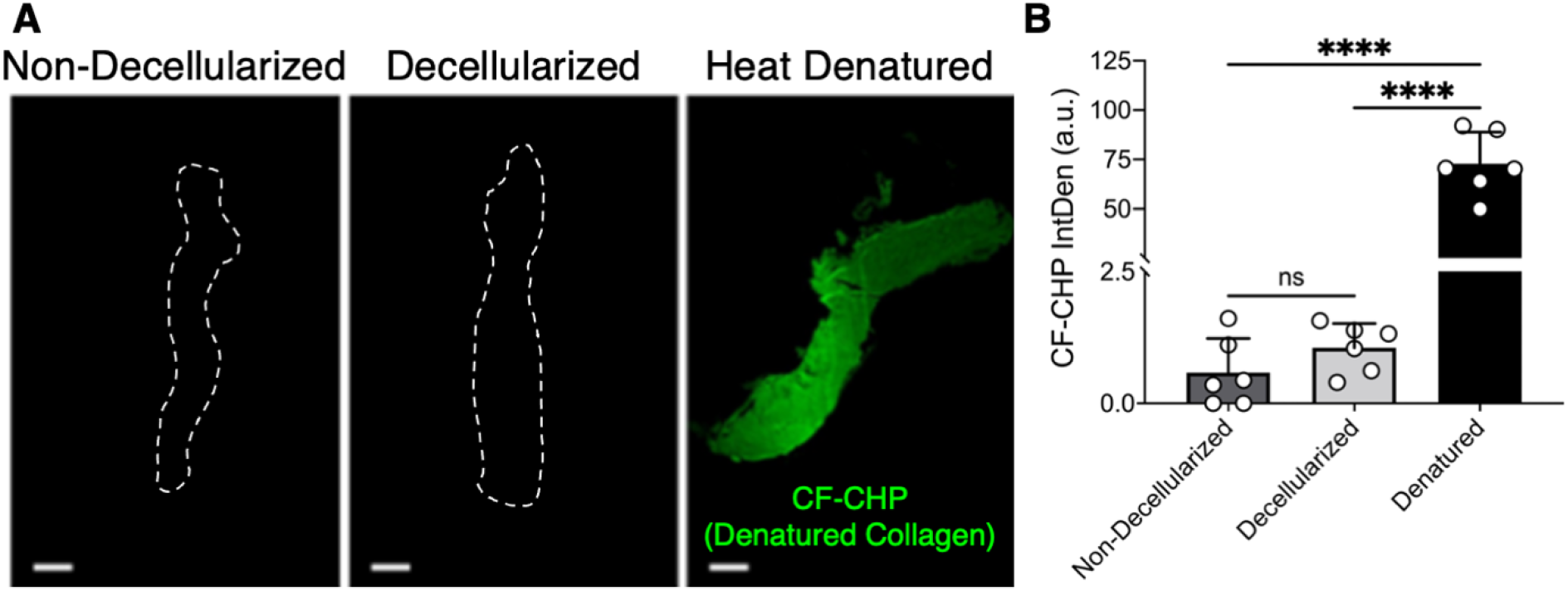
Molecular assessment of collagen denaturation using CF-CHP staining. (**A**) Representative images of CF-CHP (green indicative of denatured collagen) stained cryosections from B6 naïve patellar tendon for non-decellularized, decellularized, and heat denatured conditions. Dashed white lines indicate the tissue outline. Scale bar, 200 µm. *N* = 6 samples per group. (**B**) Quantification of CF-CHP integrated density illustrated minimal collagen denaturation in non-decellularized and decellularized tendons. Data is presented as mean ± standard deviation. \*\*\*\**P* < 0.0001.

### Bulk tissue mechanical properties were not affected by detergent-free decellularization

All tested samples failed at the tendon midsubstance. Supporting our hypothesis, there were no differences in ultimate load, stiffness, or stress relaxation between non-decellularized and decellularized B6 naïve patellar tendons (**Fig. 3A-C**). Taken together with our histological assessment, our visual observations of minimally disrupted collagen organization with decellularization are supported by these unchanged tissue-level mechanical properties. Surprisingly, while sGAGs are known to be hydrophilic macromolecules that bind water and therefore impart the viscoelastic behavior of tendon^38,39^, the percentage stress relaxation remain unchanged despite a 43.3% reduction in sGAG content.

**Figure 3.**
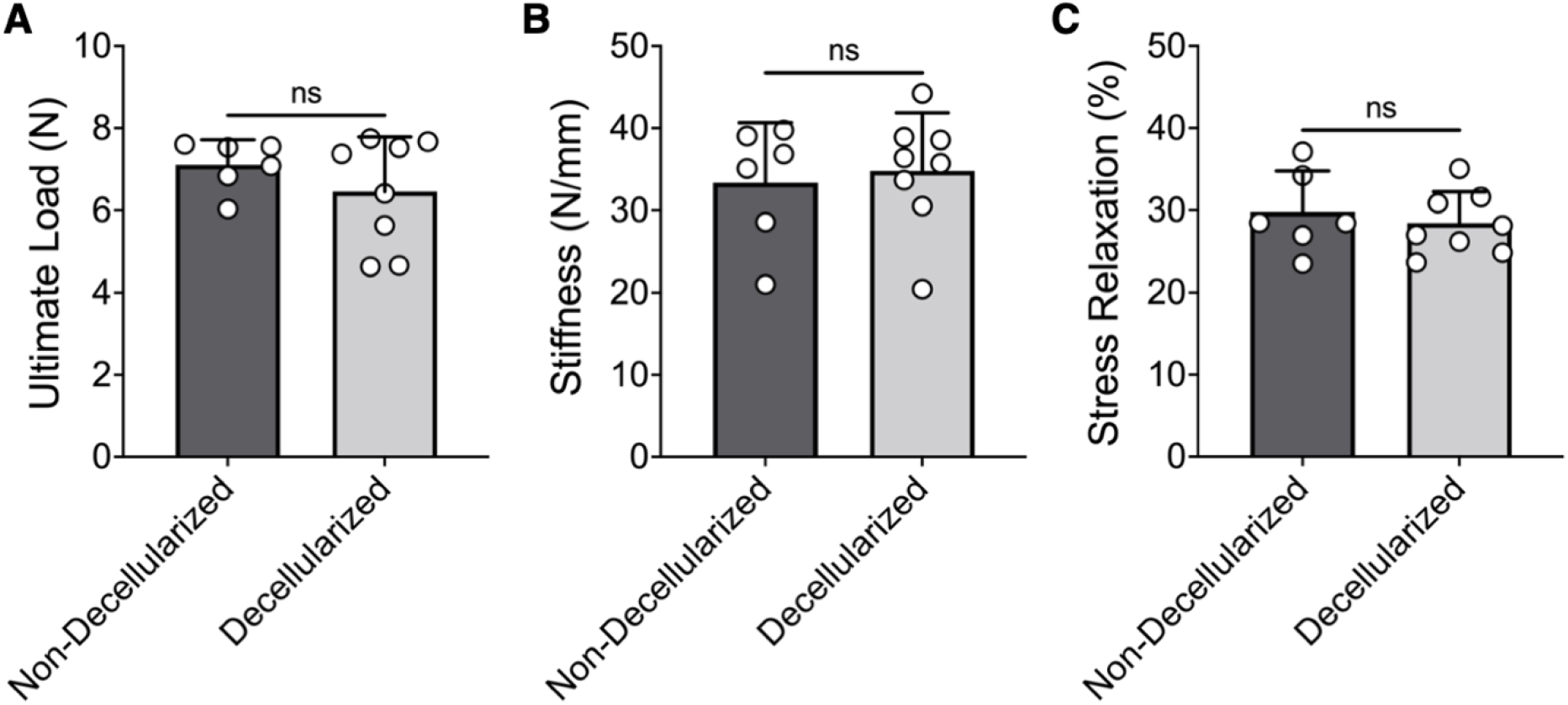
Bulk tensile testing of non-decellularized and decellularized B6 naïve patellar tendons. (**A**) Ultimate load, (**B**) stiffness, and (**C**) stress relaxation of B6 naïve patellar tendons were not affected by detergent-free decellularization. *N* = 6-8 per group. Data is presented as mean ± standard deviation.

### Tendon ECM-derived coatings are cytocompatible and promote cell survival

After 3 days under 1% FBS/serum-deprived conditions, B6 patellar tendon cells cultured on B6 naïve (*P* = 0.028) and provisional (*P* = 0.045) ECM-coated substrates exhibited significantly greater viability compared to the uncoated control (**Fig. 4A-B**). There was no difference in the total cell number between each experimental group (data not shown), indicating that tissue-specific biochemical cues harnessed by the decelluarized tendon ECM environment promote cell survival.

**Figure 4.**
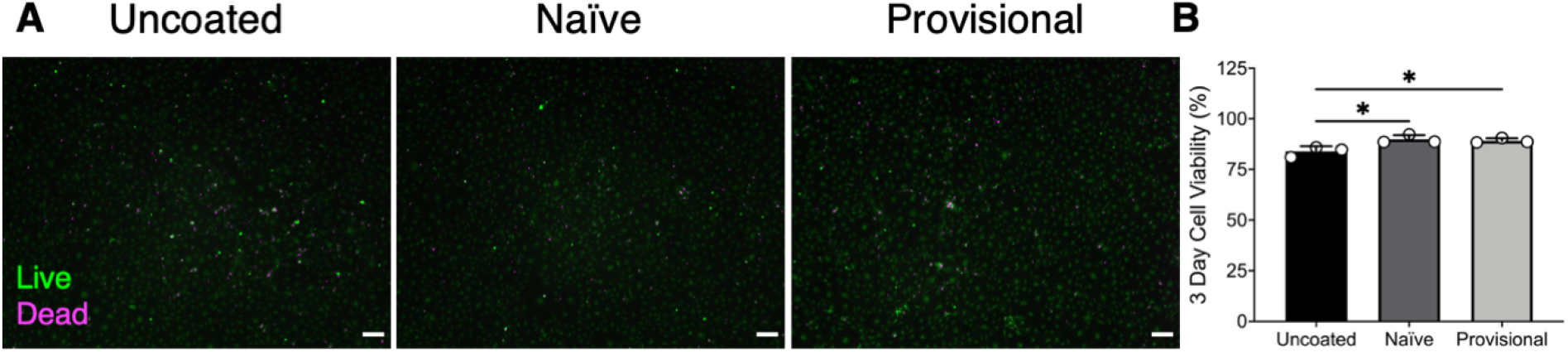
Cytocompatibility assessment of tendon cells on decellularized B6 naïve and provisional ECM-coated substrates. (**A**) Representative images of live-dead (green indicates live, magenta indicates dead) cell staining of B6 patellar tendon cells cultured on uncoated, naïve ECM-coated, and provisional ECM-coating substrates for 3 days under serum-deprived conditions. Scale bar, 200 µm. (**B**) Quantification of live-dead cell staining showed significantly increased cell viability on tendon ECM-coated substrates compared to the uncoated control. *N* = 3 per group. Data is presented as mean ± standard deviation. \**P* < 0.05.

### Single-cell morphology assessment in vitro

To further validate the utility of our detergent-free decellularization method in preserving the unique tendon ECM composition of other mouse strains, we analyzed the morphology of B6 patellar tendon cells cultured on naïve and provisional ECM-coated substrates derived from tendons of immunodeficient NSG mice^40,41^ (**Fig. 5A**). As expected, B6 tendon cells showed significantly lower spreading area (*P* = 0.017) on NSG provisional ECM-coated substrates (3879.24 ± 2134.61 µm^2^) as compared to naïve ECM-coated substrates (5052.16 ± 3532.0569 µm^2^) (**Fig. 5B**). There were no differences in cell perimeter or circularity for B6 tendon cells cultured on naïve and provisional ECM-coated substrates. Altogether, these data indicate the capacity of this detergent-free decellularization approach to evaluate differences in biologically active ECM-associated constituents (e.g., diminished pro-inflammatory cytokine signaling during NSG tendon healing) modulate tendon cell behavior.

**Figure 5.**
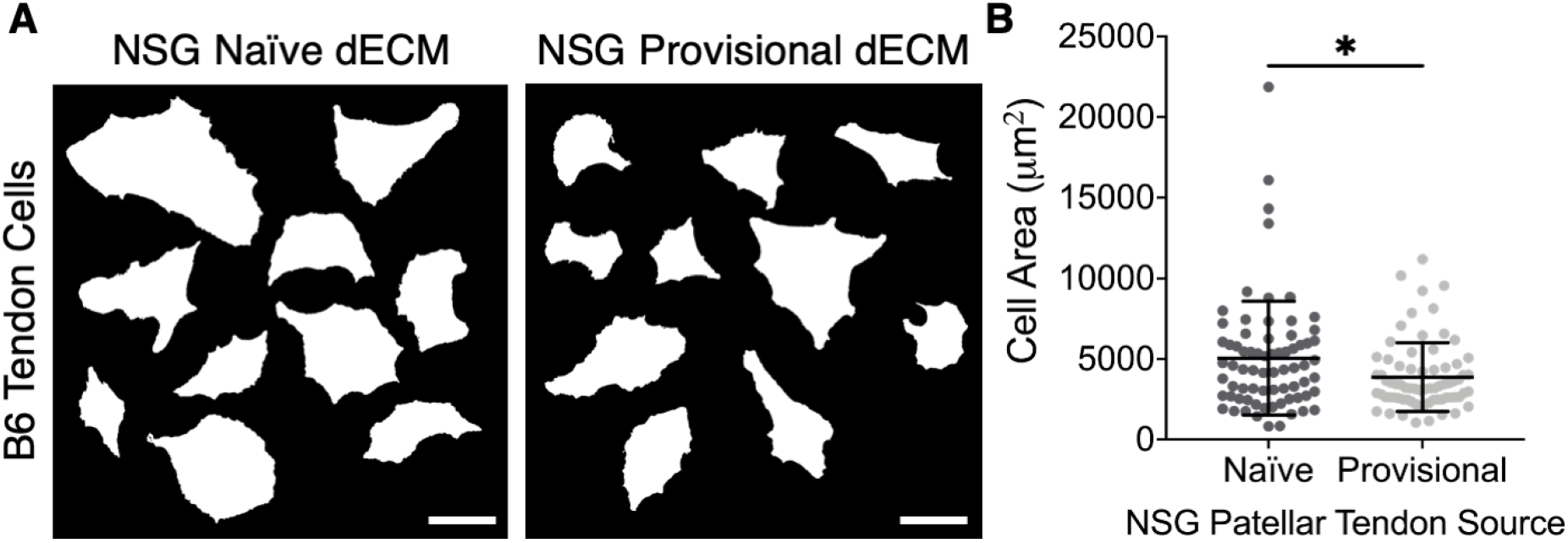
Single-cell morphological analysis of B6 patellar tendon cells cultured on NSG naïve and provisional ECM-coated substrates. (**A**) Representative cytoskeletal outlines of 10 representative cells cultured on NSG naïve and provisional ECM-coated substrates. Scale bar, 200 µm. (**B**) Quantification of single-cell morphology revealed that B6 tendon cells exhibited significantly lower spreading area when cultured on NSG provisional ECM-coated substrates compared to naïve ECM-coated substrates. *N* = 72 cells per group from 3 independent experiments. Data is presented as mean ± standard deviation. \**P* < 0.05

## Discussion

To date, the primary challenge in maximizing the potential of decellularized tissues and organs for clinical and mechanistic studies has been balancing the removal of cellular materials (i.e., as to reduce immunogenicity) and adequately preserving the complex structural and biochemical milieu of the ECM environment. The negative consequences of traditional detergent-based decellularization methods such as SDS and Triton X-100 on tendon structure and composition have been well-established. Mechanical overload of rat tail tendon fascicles has also been shown to lead to unfolding of collagen molecules as detected by CF-CHP staining^42,43^, suggesting that irreversible ECM damage achieved by either mechanical or proteolytic mechanisms may detrimentally alter cell-ECM interactions. Indeed, Veres *et al*. reported decreased accumulation of macrophage-like cells cultured *in vitro* at sites of mechanically-induced plastic deformation in decellularized collagen fibrils and an activated phenotype associated with increased cell spreading and upregulated catabolic activity.^44^ Collectively, there is mounting evidence that tissues decellularized with detergents pose a risk of compromising biological functionality and stimulating an aberrant cellular response.

Here, we successfully developed a detergent-free decellularization method for murine tendons that circumvents these concerns associated with traditional approaches. We have previously applied this method to investigate the cellular and molecular processes underlying regenerative tendon healing by isolating decellularizing provisional tendon ECM^29,30^ from the super-healer Murphy Roths Large (MRL/MpJ) mouse strain.^29,45–48^ Complementing our collagen molecular integrity data in the present study, we have previously shown that innate differences in the ECM-sequestered growth factor signaling of B6 and MRL/MpJ naïve and provisional tendons are retained after decellularization.^29^ We also previously demonstrated that B6 tendon cells cultured on decellularized MRL/MpJ dECM-coated substrates exhibited enhanced cell proliferation, elongation, and the formation of cellular protrusions that are characteristic of MRL/MpJ tendon cell behavior.^29^ Excitingly, these findings validate the versatility of our detergent-free decellularization method and its untapped capacity to mechanistically interrogate tendon ECM biology using other transgenic mouse strains, sex-based comparisons, enzyme-mediated ECM depletion (e.g., elastase for elastin or plasmin for fibrin), and other models of tendon injury such as chronic overuse through strenuous exercise or fatigue loading.^49^

Although the decellularized B6 provisional ECM exceeded the 50 ng dsDNA/per mg of dry weight threshold proposed by Crapo *et al*.^10^, we have previously implanted MRL/MpJ dECM-derived hydrogels into injured B6 patellar tendons *in vivo* and improved the tissue structure and mechanical properties without observing a deleterious inflammatory reaction indicative of an adverse immunogenic response.^30^ Thus, our study opens up the avenue for future studies to explore whole-tissue recellularization strategies through surgical repair [e.g., anterior cruciate ligament (ACL) reconstruction], *in vivo* transplantation^50^, or e*x vivo* bioreactor and organ culture systems to assess the crosstalk between ECM and mechanical regulation.

## Acknowledgments

The authors thank Claudia Fischbach-Teschl for providing training and technical assistance with their epifluorescence microscope.

## Disclosure Statement

No competing financial interests exist.

## Funding Information

This work was supported by the National Institutes of Health (NIH) under the following award numbers: R01AR608301 (to N.A.P.), R01AR052743 (to N.A.P.), and S10RR025502 (to Cornell Biotechnology Resource Center). The authors also acknowledge support from the National Science Foundation (NSF) Graduate Research Fellowship Program (GRFP) DGE-1650441 (to J.C.M.).

## Notes

### Competing Interest Statement

The authors have declared no competing interest.

